# Novelty by Furcation and Fusion: How tree-like is evolution?

**DOI:** 10.1101/113613

**Authors:** Todd H. Oakley

## Abstract

Novelty and innovation are fundamental yet relatively understudied concepts in evolution. We may study novelty phylogenetically, with a key question of whether evolution occurs by tree-like branching, or through exchange of distantly related parts in processes akin to horizontal transfer. Here, I argue that except at the lowest levels of biological organization, evolution is not usually tree-like. Perfectly vertical inheritance, an assumption of evolutionary trees, requires simultaneous co-duplication of all the parts of a duplicating or speciating (which I collectively call 'furcating') biological feature. However, simultaneous co-duplication of many parts usually requires variational processes that are rare. Therefore, instead of being perfectly tree-like, evolution often involves events that incorporate or fuse more distantly related parts into new units during evolution, which herein I call 'fusion'. Exon shuffling, horizontal gene transfer, introgression and co-option are such fusion processes at different levels of organization. In addition to co-duplication, units under phylogenetic study must individuate (gaining evolutionary independence) before they can diverge. A lack of individuation erases evolutionary history, and provides another challenge to tree-like evolution. In particular, biological units in the same organism that are the products of development always share the same genome, perhaps making full individuation difficult. The ubiquity of processes that fuse distantly related parts or oppose individuation has wide ranging implications for the study of macroevolution. For one, the central metaphor of a tree of life will often be violated, to the point where we may need a different metaphor, such as economic public goods, or a ‘web of life’. Secondly, we may need to expand current models. For example, even under the prevailing model of cell-type evolution, the sister-cell-type model, a lack of complete individuation and evolution by co-option will often be involved in forming new cell-types. Finally, these processes highlight a need for an expansive toolkit for studying evolutionary history. Multivariate methods are particularly critical to discover co-variation, the hallmark of an absence of complete individuation. In addition to studying phylogenetic trees, we may often need to analyze and visualize phylogenetic networks. Even though furcation - the splitting and individuation of biological features - does happen, fusion of distant events is just as critical for the evolution of novelties, and must formally be incorporated into the metaphors, models, and visualization of evolutionary history.

## I. Introduction

To understand how biodiversity and complexity arise, we must understand the evolution of new things - so called biological novelties (structures) and innovations (functions) (Wagner, 2015). Biodiversity may be defined as variety of species and complexity defined as variety of parts (McShea and Brandon, 2010). Greater variety requires novelty and/or innovation. We have much to learn about the evolution of novelty and innovation, and their relationship to complexity, diversity, and disparity (McShea and Brandon, 2010; Wagner, 2014). Every species, every trait, and every gene of every species is the evolutionary product of novelties at some point in the past. Without novelty, evolution could not occur. Although novelties occur in populations, complexity and biodiversity are macroevolutionary topics, whose evolution must be studied phylogenetically. Complex features like eyes and species and photosynthesis did not originate in a single population. Instead, complex biological entities are composites of multiple novelties that happened across time, in multiple different populations/generations. An important part of understanding evolution is to study the timing, environmental context, and order of these macroevolutionary events. For example, what are the novelties that led to the evolution of complex features, like eyes, feathers, flowers, or metabolic pathways? When and how did the novelties comprising these features come together, and what was the environment like at the time (Oakley and Speiser, 2015)?

In order to facilitate discussing these topics, I will herein make two distinctions. First, I consider that biological novelties evolve in two different ways, which I call furcation (duplication-like processes) and fusion (merging formerly separate parts), which result in distinguishable historical patterns when studied phylogenetically. These patterns boil down to asking how tree-like is evolution. Furcation is tree-like, fusion is network-like. Second, I will distinguish genetic and developmental systems. Genetic systems are passed from generation to generation, and include domains, genes, and genomes (species). Developmental systems develop anew in each generation (Musser and Wagner, 2015; Wagner, 2014), and include morphology, organs, and cell-types. The first distinction (furcation versus fusion) is important for my discussion because phylogenetic methods used to study novelty, tended to assume one process (furcation), and exclude the other (fusion). The second distinction (genetic versus developmental systems) is important because the phylogenetic study of developmental systems is far less mature than the study of genetic systems, and the latter may have a lot to teach the former.

The first process to produce novelties involves duplication, individuation (which means the copies gain the ability to differentiate), and divergence - which Oakley et al (2007) termed ‘furcation’. You might be more comfortable with the terms ‘duplication’ and ‘speciation’ - which are certainly useful - but furcation as a term has some advantages. First, furcation is similar to ‘duplication’, but more general in that it applies to any level of biological organization, including gene and species levels. Second, ‘duplication’ does not necessarily include individuation, which furcation explicitly does. Finally, furcation applies to both copying and splitting, and includes both duplication and multiplication. A familiar example of furcation is gene duplication. Within a genome, one of multiple mechanisms copies a gene, allowing the genes to diverge. For many genes, simply copying them to different places in the genome allows individuation (unless concerted evolution (reviewed in Liao, 1999) occurs). The result of gene furcation is a biological novelty because there are now twice as many genes. This furcation also increases the structural complexity of the genome because it now has a greater variety of parts (McShea and Brandon, 2010). Besides gene duplication, furcation also includes speciation, vicariance, language differentiation, and the evolutionary division of a cell-type or organ into descendent cell-types or organs.

A second process producing novelties is copying to make new combinations, also called ‘fusion’ (Oakley and Rivera, 2008) because multiple distantly related parts fuse to form a new unit. Like furcation, fusion also happens at all levels of biological organization (Table 1). Herein, I use fusion to encompass processes like horizontal transfer, hybridization, introgression, creolization (hybridization of languages), co-option, and exon shuffling. The latter, exon shuffling, provides a rather straightforward example. Within a genome, one of multiple mechanisms copies part of a gene (e.g. a protein domain), inserting the copy next to an unrelated domain to form a new gene. A specific example is *Pax-6,* which contains a *Paired* and *Homeobox* domain. The first *Pax-6* gene was a novelty, originated by fusion of unrelated domains. It increased the structural complexity of that fateful ancestor’s genome because it gained a new and different gene. Figure 1 presents a generalized cartoon of furcation (A) and fusion (B). I note that these processes are inter-dependent because fusion involves furcation of the lower level units. In the *Pax-6* example, we can create perfectly branching phylogenetic trees of the domains themselves. It is only at the higher level of the gene where we see the fusion of the distantly related *Paired* and *Homeodomains*.

**Figure 1.**
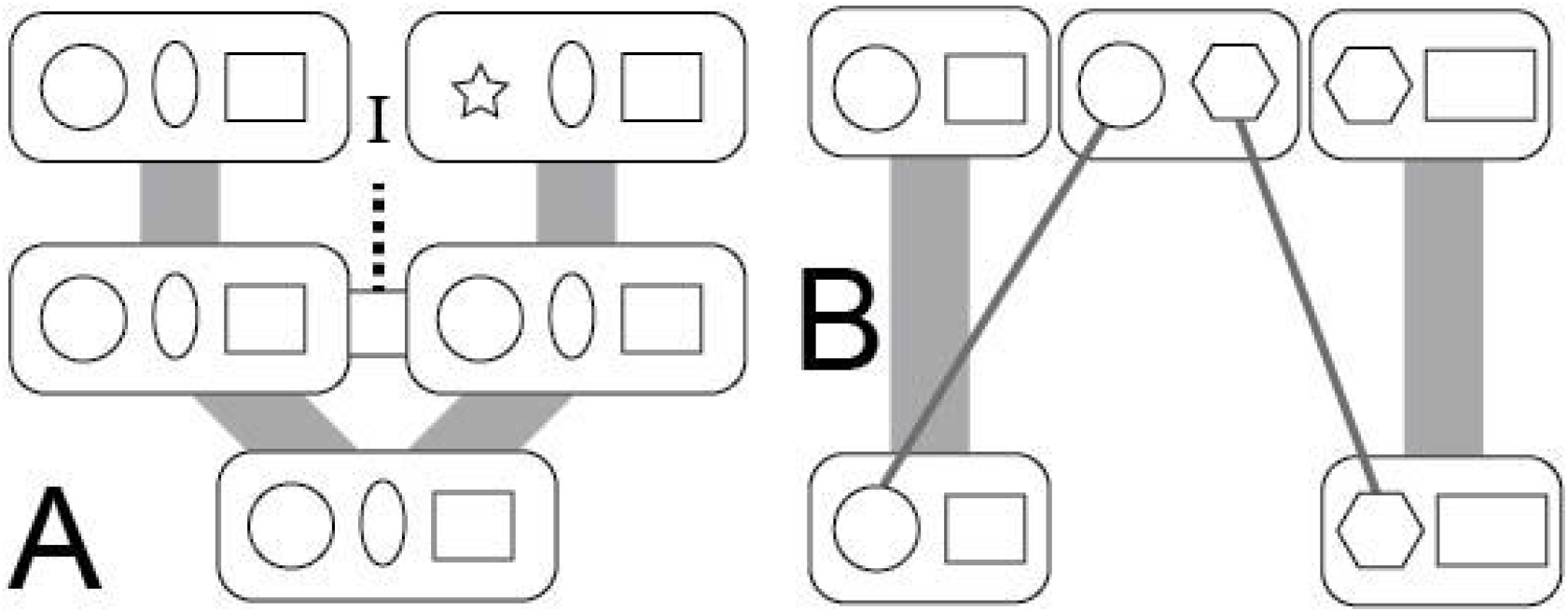
Novelties originate by A. Furcation or B. Fusion. The rounded rectangles represent a higher-level biological unit and the shapes inside are parts of that unit. For example, the figures could represent domains in a gene or genes in a species. Ancestral biological units are at the bottom, descendant biological units toward the top. Ancestor-descendent relationships are depicted by grey lines. First, a process copies the unit - a gene is duplicated or populations establish. Furcation involves copying, individuation (I), and divergence. The horizontal bar (h) represents a homogenizing process. For genes, this is concerted evolution, for populations, it may be gene flow. Next, the units become individuated, losing the homogenizing process. For populations, this could be reproductive isolation. Finally, the units diverge from each other (notice the ‘star’ shape instead of the circle). For genes and species this could be through the fixation of different mutations. Fusion involves the assembly of copied parts into a new biological unit. Again, rounded rectangles are a higher level biological unit, with shapes representing parts. The units could be cell-types or genes, and the shapes would then be expressed genes or domains, respectively. During fusion, parts are copied to form a novel combination. In this diagram, the fused parts are the same in number (one part from each source). However, one part from one unit could be added with many parts of another. An example would be horizontal transfer, where one genome adds a single gene from an unrelated genome.

**Table 1.**
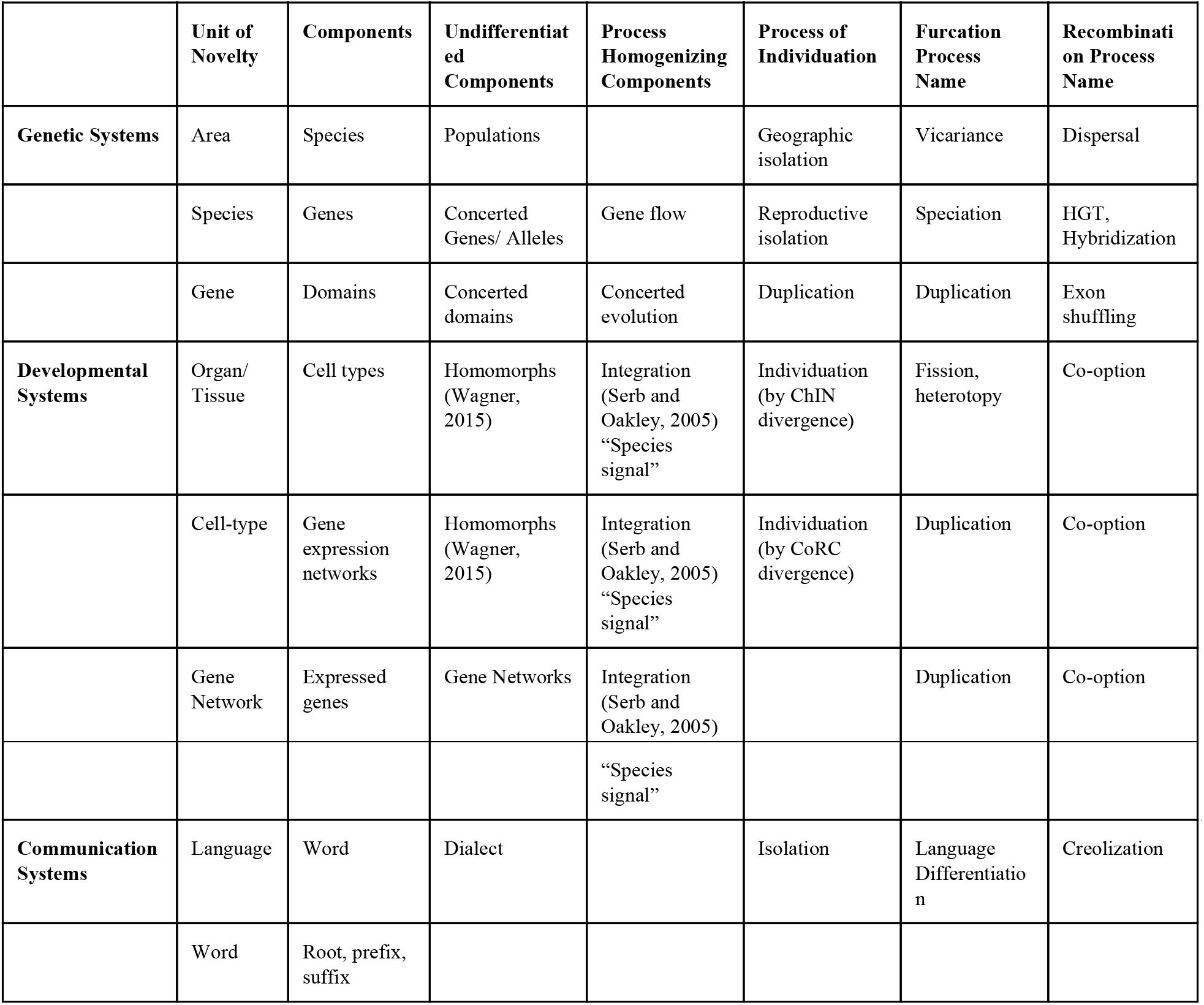
Novelty concepts across levels of organization

|  | Unit of Novelty | Components | Undifferentiated Components | Process Homogenizing Components | Process of Individuation | Furcation Process Name | Recombination Process Name |
| --- | --- | --- | --- | --- | --- | --- | --- |
| Genetic Systems | Area | Species | Populations |  | Geographic isolation | Vicariance | Dispersal |
|  | Species | Genes | Concerted Genes/ Alleles | Gene flow | Reproductive isolation | Speciation | HGT, Hybridization |
|  | Gene | Domains | Concerted domains | Concerted evolution | Duplication | Duplication | Exon shuffling |
| Developmental Systems | Organ/ Tissue | Cell types | Homomorphs (Wagner, 2015) | Integration (Serb and Oakley, 2005) “Species signal” | Individuation (by ChIN divergence) | Fission, heterotopy | Co-option |
|  | Cell-type | Gene expression networks | Homomorphs (Wagner, 2015) | Integration (Serb and Oakley, 2005) “Species signal” | Individuation (by CoRC divergence) | Duplication | Co-option |
|  | Gene Network | Expressed genes | Gene Networks | Integration (Serb and Oakley, 2005) |  | Duplication | Co-option |
|  |  |  |  | "Species signal" |  |  |  |
| Communication Systems | Language | Word | Dialect |  | Isolation | Language Differentiation | Creolization |
|  | Word | Root, prefix, suffix |  |  |  |  |  |

Herein, I will also distinguish between genetic and developmental systems. I use the term ‘genetic systems’ to include protein domains, chromosomes, genes, and species. The application of phylogenetic and historical methods to genetic systems is familiar. When we study the evolutionary history of domains and genes, we can study those structures directly. Changes in gene sequence evolve over time such that similarity in genes should usually decrease over evolutionary time. I admit, species are a bit different, but I include them here with genetic systems in part because we sometimes equate gene trees directly with species trees. But mainly, I include species within genetic systems because studying their phylogeny is very familiar, and I want to focus most on phylogenetic thinking in developmental systems, which is less familiar. I use ‘developmental systems’ to include protein interaction networks, cell types, organs, tissues, morphological traits, behaviors, and languages. Developmental systems do not have continuity between generations, but develop anew in each individual (Wagner, 2014). Although less familiar, a small set of recent publications has gone beyond species and gene trees, to study the phylogenetics of these developmental systems (Arendt, 2008; Geeta, 2003; Kin, 2015; Musser and Wagner, 2015; Oakley, 2003; Oakley et al., 2007; Serb and Oakley, 2005).

Herein I argue that phylogenetic methods make strict assumptions of co-duplication and individuation that are commonly violated at every level of biological organization. After explaining examples from genetic systems, I discuss developmental systems. Given that phylogenetic trees assume only vertical inheritance, I then explore a previous proposal to change the central metaphor of macroevolution - the tree of life - which discourages our thinking about horizontal, fusion processes. In particular, I explore what thinking about evolution from an economic goods perspective (McInerney et al., 2011) might add. I conclude with a brief review of models and visualization of phylogenetic networks, which are required when strict branching processes are violated.

## II. Evolutionary trees assume co-duplication and individuation

Because evolutionary trees assume ancestor-descendent inheritance, they assume all components duplicate and individuate simultaneously. A focus on using gene trees to infer species trees makes the assumption of co-duplication easy to understand. The goal of species trees is to depict common ancestry of the species, with each node of the tree indicating speciation of a hypothetical common ancestor into descendent species. Phylogeneticists usually assume that all genes belonging to a species’ genome are inherited from a direct ancestor, without horizontal gene transfer. For this assumption to hold, all genes of each ancestor must have split simultaneously at each speciation event. If so, all gene trees are congruent with the species tree. What process could cause all genes of a genome to split concurrently into two genomes?

A simplistic model of speciation could cause all genes of an ancestral genome to split and individuate simultaneously. Imagine two populations of an ancestral species become separated by a geographic barrier that fully prevents gene flow between the populations. In this simple example, the ancestral genome splits into two descendent genomes: one genome on either side of the geographic barrier. Without gene flow, the descendent genomes must be fully individuated because they share no information with each other. In this case, every gene of the ancestral genome is ‘duplicated’ (geographically, each within a separate population) and immediately individuated. Individuation is clear in the absence of gene flow because the copies of each gene in the descendant genomes now evolve independently of each other.

We make similar assumptions when generating phylogenetic trees at other levels of biological organization. For example, genes are usually composed of components like protein domains. Therefore, a phylogeny of a gene family assumes that all domains within the gene family duplicated and individuated simultaneously. In other words, if we were to create separate trees of those domains, we would assume those trees would be identical, perfectly congruent with each other. For example, we expect the *Paired* and *Homeobox* domains of *Pax-6* shared their evolutionary history through speciation and duplication events, such that phylogenetic analyses of each separate domain would yield congruent results. Even though these assumptions of phylogenetic trees are strict and common, we know they are often violated.

## III. Co-duplication and individuation are often violated in genetic systems

Unfortunately for phylogenetics, many evolutionary processes violate the assumptions of vertical inheritance. In the evolution of populations and genomes, multiple well established processes like introgression, and horizontal gene transfer violate strict vertical inheritance, violate co-duplication, and cause incongruence between gene trees and species tree (Hahn and Nakhleh, 2016; Maddison, 1997; Mallet et al., 2016). Similarly, in the evolution of genes, exon shuffling often fuses distantly related domains together to form new genes. Sticking with our *Pax-6* example, even though *Paired* and *Homeobox* domains have had congruent histories since they joined forces at the origin of *Pax-6,* each domain also has a deeper history that is separate from each other (Oakley and Rivera, 2008). As in *Pax-6,* multi-domain proteins are enormously common, such that fusion of distantly related or unrelated domains must be an important mode of gene origin (Haggerty et al., 2013).

Besides fusion of distant domains or genes, a lack of individuation also violates assumptions of phylogenetic trees. In order for replicated parts to diverge, they must “individuate”; they must have the ability to evolve differences. A lack of individuation keeps duplicated parts the same such that they cannot track evolutionary history. A lack of individuation, then, is another way the assumptions of phylogenetics are violated. At the level of genes, concerted evolution is a process within a genome leading to homogenization of duplicates. As a result, old paralogs are more similar within a species than they are to orthologs in a different species. This is not what we might at first expect. Instead, once a gene is duplicated, we expect the copies to accumulate mutations separately, differentiating from each other more and more over time. McShea and Brandon (2010) call this idea the Zero Force Evolutionary Law (ZFEL): evolutionary entities should diverge unless counteracted. In this case, instead of the duplicate genes evolving individually, they could evolve in concert, a result of homogenizing mutational mechanisms like unequal crossing over, which could be influenced by natural selection. Therefore, concerted evolution may be conceived as a lack of individuation. As a result, genetic variety and complexity do not increase for duplicated genes that undergo concerted evolution. Concerted evolution erases historical signal, so the relationships of the genes do not represent the historical pattern of replication.

## IV. Developmental systems and tree-like evolution

Driven by inexpensive and nearly comprehensive expression data for organs, tissues, and individual cells, evolutionists now have the ability to study novelty and its role in the complexity of developmental systems. Which is more important in the origins of developmental systems - furcation or fusion? In other words, to what extent are the evolutionary histories of developmental systems tree-like? Here, I argue that for two primary reasons, the assumptions of phylogenetics will very often be violated during the evolution of developmental systems. First, the shared genome of developmental systems makes individuation more difficult, leading to process akin to concerted evolution that erase historical signal. Second, I argue that variation in developmental systems will often involve co-option, so the evolution of developmental systems is a network, not a tree.

### a. A shared genome challenges individuation

Individuation occurs when replicated biological entities lose integration and diverge from each other. However, sometimes replicated features do not individuate, violating the expectation of divergence assumed in tree-like evolution. As I discussed above, genes may evolve by concerted evolution. Similarly, we may view gene flow between populations as opposing individuation because alleles may be recombined between those populations. Hybridization and introgression also result from imperfect reproductive isolation, causing genomes to evolve not solely by splitting and divergence, but by incorporating genes from more distant relatives. Only with complete individuation does evolution meet the assumptions of phylogenetic trees.

In addition to these genetic systems, developmental systems may also evolve in concert. Serial homologs often remain variationally integrated after they duplicate, a process called morphological or developmental integration (Cheverud, 1996; “Morphological Integration and Developmental Modularity,” 2008; Zelditch and Fink, 1995). Serb and Oakley (2005) hypothesized (see their Fig 2, middle column bottom row) that cell types could be homogenized by a process they termed “developmental integration”. Interestingly, some data now indicate that developmental integration of tissues and cell types happens during evolution, although the specific underlying mechanism remains unknown. Recent authors term developmental integration “the species signal” (Liang et al., 2016; Musser and Wagner, 2015) or use the term “concerted evolution” for any level of biological organization, pointing out developmental integration in several data sets. For example, cephalopod organs are more similar in gene expression within a species than the similarity of organs in gene expression between species (Pankey et al., 2014). Liang et al (2016) provide a statistic to quantify developmental integration and the authors find more integration between developmentally related tissues.

**Figure 2.**
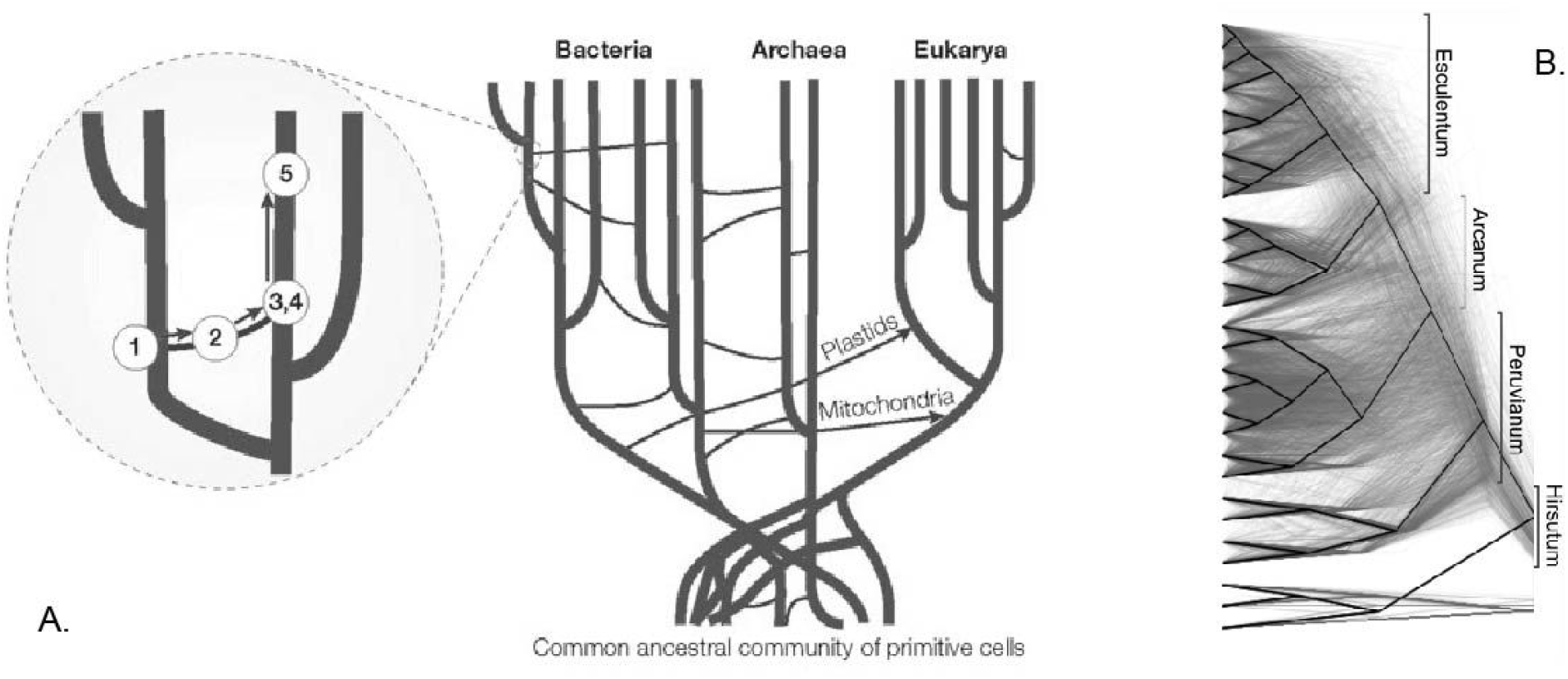
Genomes often incorporate distantly related genes. **A**. Horizontal Gene Transfer is a common process, especially in prokaryotes and in a common ancestral community of primitive cells. Genes or entire genomes (plastids) are ‘fused’ with a distantly related genome. Figure from (Smets and Barkay, 2005). **B**. Gene trees (thin lines looking like a grey cloud) are often incongruent with species phylogeny (black branches) due to processes like introgression, here illustrated in tomato plants. Figure from (Pease et al., 2016).

**Figure 3.**
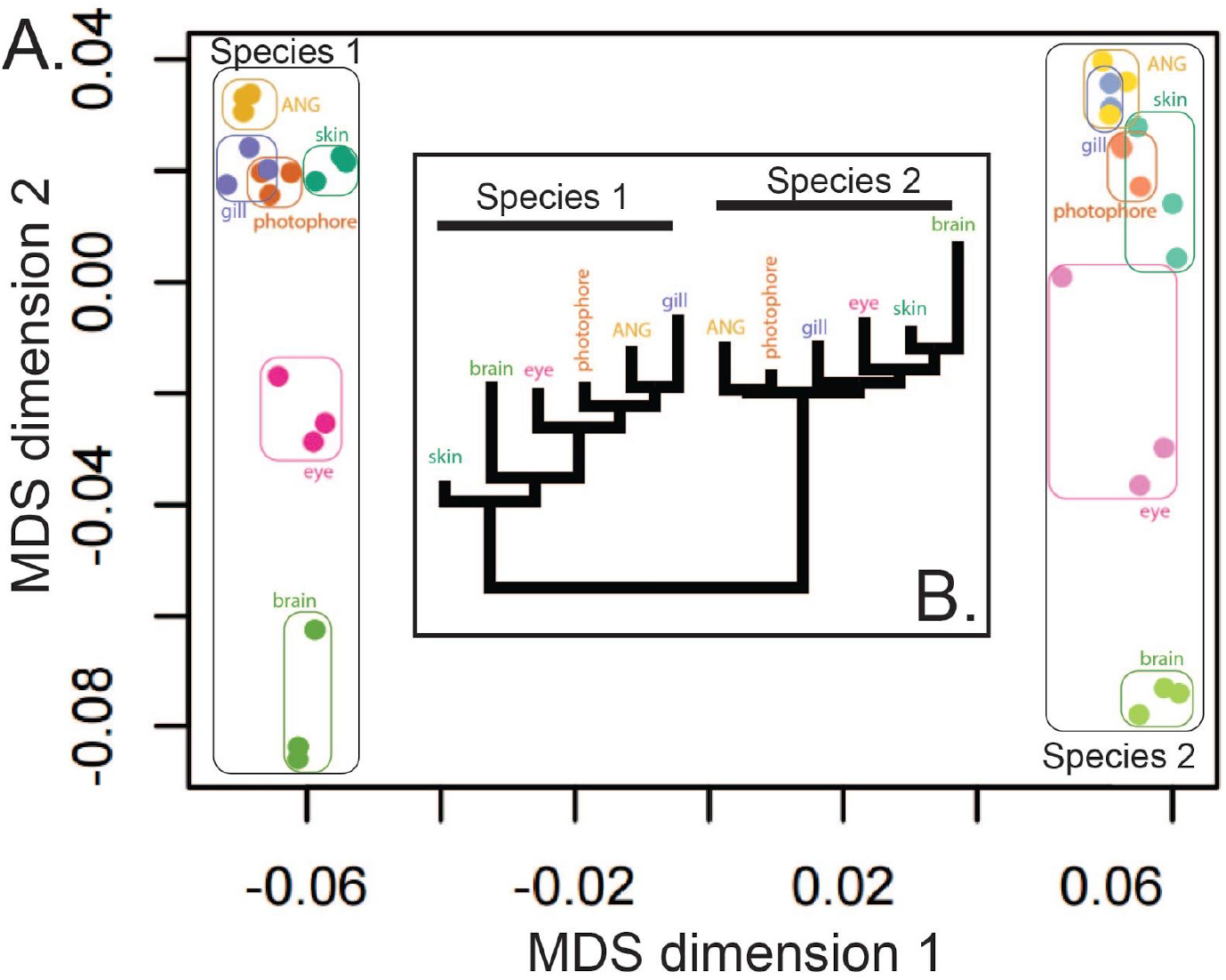
Overall gene expression of cephalopod organs clusters strongly by species. **A.** Multi-Dimensional Scaling (MDS) analysis strongly differentiates organs from two cephalopod species in dimension 1, even for homologous organs. Analysis from (Pankey et al., 2014). **B**. A phylogenetic tree from the same cephalopod gene expression data groups all organs from each species into reciprocally monophyletic groups. Without developmental integration, we would expect homologous organs from each species to group together. One interpretation of the observed result is that developmental integration (concerted evolution) is driving co-variation of expression within species.

Perhaps the shared genome of developmental systems makes complete individuation difficult or even impossible. In fact, it seems likely there is no analog of complete ‘reproductive isolation’ for the cells and organs of a multicellular organism. The concept of ‘housekeeping genes’ provides an intuitive reason why. In principle, housekeeping genes each have one instance in a genome, they are expressed in all cells, and they are required for foundational cellular processes. A change in the genomic sequence of a housekeeping gene then occurs in all cells where the protein is expressed, no matter how distantly related those cells are from each other on a cell-type phylogeny. Even though housekeeping genes may have interactions with other proteins that are specific to tissue-types, broadly expressed proteins also have the most protein-protein interactions (Bossi and Lehner, 2009), such that a change in protein sequence will likely lead to integrated changes in function across cell and tissue types. Individuation of cell types and organs may also be difficult with respect to gene expression. A gene expressed in many cell types, perhaps even distantly related cell types, may regulate expression through the same specific genomic changes. From this perspective, cell types may not individuate because the same genomic changes, perhaps to a transcription factor, could cause concerted change to replicated cell types.

### b. Co-duplication, co-option, and variation of developmental systems

Tree-like evolution of biological entities implies that all components of the entity replicate and individuate simultaneously, which I call furcation. Furcation requires a very specific variational process. As I already pointed out for species, perfect furcation requires populations to split without any subsequent gene flow, allowing all genes of a genome to divide and differentiate at one specific point in time. For gene trees to be tree-like, I also mentioned that the entire gene must duplicate at once, not just a fraction of that gene’s domains. Such modes of variation in these genetic systems are conceivable. For the origin of genes especially, co-duplication of all domains is very common. How commonly do we expect co-duplication of the parts of developmental systems?

At least to a first approximation, organs and cell-types seem to replicate, but do they undergo perfect furcation so that we can study them with phylogenetic trees? Or is fusion (co-option) so common that we must use evolutionary networks to visualize the histories of developmental systems? Developmental systems are comprised of many components, mainly expressed genes. To meet the strict assumptions of a phylogenetic tree for developmental systems would require that all expressed genes be duplicated simultaneously at the time of origin. Oakley et al. (2007) rejected co-duplication for multiple arthropod photoreceptor systems. For example, even where duplicated opsin genes are expressed in potentially furcated arthropod eyes, the opsin duplication was much younger than the origin of the eyes themselves (Oakley et al., 2007). Should we ever expect all the genes used in a complex developmental feature to duplicate simultaneously? The possible origin of such variation seems unlikely. In order for all genes of - say - an eye to duplicate would require full genome duplication, or all the genes to be syntenic and duplicated as a block. Full genome duplication is quite plausible, and even implicated in the origin of duplicated phototransduction networks of vertebrate rods versus cones (Nordström et al., 2004; Oakley and Plachetzki, 2011; Plachetzki and Oakley, 2007). But while operons of co-regulated genes are common in prokaryotes and blocks of genes are often horizontally transferred together as ‘caravans’ in eukaryotes (Wisecaver and Rokas, 2015), the syntenic organization of co-functioning genes does not seem to be a general feature of animal genomes. Therefore, strict furcation of tissues and organs would seem to be rare. Without origination of a new organ by duplication of an ancestral organ and all its genes, organs are necessarily evolving by fusing together (co-opting) genes used in other contexts.

What if we relax this requirement of duplication of gene structure and consider gene expression into a new place as duplication of function? For example, ectopic expression of the specification genes of an eye could lead to organ development and gene expression in a new place. The eye genes themselves are not duplicated, but expression is replicated in a new place. We might think of these specification genes as the Character Identity Network (ChIN) described by Wagner (Wagner, 2014, 2007). I argue that by only replicating expression, changes to the genomic sequence of those genes will cause concerted changes where those genes are expressed, in both the original and ectopic eye. In other words, the replicated eyes cannot be fully individuated. For this reason, we may expect to commonly encounter developmental integration across organ and tissue types. As in concerted evolution of genes, the history of developmentally integrated organs will be erased. As a result, integrated organs will be each others closest relatives, even if they probably separated long ago. Some empirical data sets support this idea (Liang et al., 2016; Musser and Wagner, 2015; Pankey et al., 2014). In cephalopods, traits including eyes, liver, skin group first by species in multi-dimensional plots of gene expression. If the traits were individuated, we would expect homologous organs to group together - eyes with eyes, and liver with liver. Organs and tissues are amalgamations of multiple different cell types. Might we expect a similar or different pattern at the level of cell type evolution?

For the same reasons as organ evolution, I argue that new cell-types should also usually involve co-option, thus violating strict assumptions of phylogenies. One model of cell-type evolution is the sister-cell model. The sister cell type model posits that new cell types often arise by furcation of an existing cell type into descendent cell types (Arendt et al., 2016). Cells are routinely multiplied during organismal growth, so duplication/multiplication is not the interesting bit of cell type furcation. Instead, individuation and divergence is key. How do multiplied cells diverge from each other? The sister cell type model focuses on a Core Regulatory Complex (CoRC) to define cell types, such that changes to CoRCs cause new cell types. A CoRC is a group of regulatory proteins, probably transcription factors, that determine the set of effector (or ‘downstream’) genes that are expressed in a cell type. Under what conditions could sister cell types meet the strict assumptions of phylogenies, which discount the possibility of co-option?

Just as with other developmental systems, I argue the origin of sister cell types will usually involve incomplete individuation and co-option, thus violating strict assumptions about phylogenetic trees. Just as with organs, all genes expressed in sister cell types would have to co-duplicate before complete individuation is possible. Instead, genes comprising housekeeping modules will be expressed in both sister cell types, so changes in sequence to these genes will affect both cell types. In addition, differentiation of the CoRC will often involve co-option. For example, Arendt et al. (2016) posit that CoRCs may change the set of transcription factors determining cell identity. Unless these CoRC transcription factor genes themselves duplicate and specialize for each daughter cell type, a new CoRC will involve co-option of transcription factors. Furthermore, the new transcription factors may cause expression of new modules of genes (called apomeres) in the sister cell types. Again, unless all those genes are co-duplicated and differentially expressed across sister cell types, they will be co-opted from other cells. Just as with organs, it seems unlikely that all genes expressed in a particular module will be syntenic and co-duplicated. As a result of incomplete individuation, a first step to analyzing the evolution of developmental systems may be to quantify integration of the traits (Liang et al., 2016). Next, we might study relationships of the developmental systems to each other, independent of the co-variation caused by being in the same species, perhaps by using confirmatory Factor Analysis. Because of the necessity of co-option in the origin of cell types and organs, I argue that this history should usually be analyzed as cell-type networks.

## V. Implications of fusion

The metaphor of a tree of life occupies a central place in our understanding of evolution. However, as I reviewed briefly above, fusion by horizontal transfer and hybridization violate assumptions of phylogenetic trees, causing incongruence in genetic systems between the history of genes and the species that contain those genes (Hahn and Nakhleh, 2016; Maddison, 1997). Additionally, I argued above that developmental integration and co-option may be commonplace in developmental systems, causing incongruence between the history of gene expression and the cell types and organs that express those genes. Therefore, the history of life cannot accurately be visualized as a single phylogenetic tree. Is there an alternative metaphor to the tree of life? In this section, I discuss alternatives to a strict tree of life. First, I discuss an alternative metaphor, then I mention alternative models and modes of visualization.

### a. An alternative metaphor

One alternative to strict tree-thinking is an economic goods metaphor (McInerney et al., 2011), borrowed from economics, where biological entities are ‘non-excludable’. Like the new economy (DeLong and Froomkin, 1997) of the internet age, where information is cheap to copy and can be used by anyone simultaneously, biology may employ an analogous information economy. In the context of species phylogeny, ‘non-excludable’ means that the the components of species (e.g. genetic material) can be transferred horizontally from species to distantly related species. At other levels, protein domains (the components of genes) may be transferred to distantly related genes, and cell types or organs might co-opt expression of genes in distantly related cells or organs.

Perhaps the biggest advantage of considering an economic goods metaphor in macroevolution is to force us to ease the assumption of strict vertical inheritance of biological features. Evolutionary biologists already know that vertical inheritance is not ubiquitous. Yet representing evolution as a phylogenetic tree assumes it. Even when fusion is noted, by pointing out incongruence of a component’s history with a higher level phylogeny (e.g. gene tree incongruence with species tree or domain tree incongruence with gene tree), it is easy to view incongruence as simply an exception to the rule. However, “Scientific illustrations are not frills or summaries; they are foci for modes of thought (Gould, 2010)”. From this perspective, perhaps changing our mode of thought from tree to economic goods (or perhaps more poetically to a ‘web of life’), and by often changing our illustrations from trees to networks, we could reinforce a mode of thought that considers fusion as an equal alternative to furcation. Next, I explain the economic goods metaphor, with a focus on its currently limited use in macroevolution.

#### Economic classification of goods

In economics, “goods” may be classified into a 2×2 matrix, forming four categories. In practice, the categories are usually not discrete alternatives, but instead form axes with continuous variation. One axis asks how “excludable” and the other axis asks how “rival” is a particular good. Non-excludable goods can be ubiquitously accessed, whereas excludable goods have restricted access. For example, the air we breath has low excludability because we cannot easily (ethically) prevent anyone from breathing the atmosphere. However, a theme park is quite excludable; only those with a ticket may enter. On the other axis, highly rivalrous goods are exhaustible, but non-rival goods are unlimited. A parking space is rivalrous because if I park there, no one else can. In contrast, a public radio signal is non-rivalrous. My listening to 92.9 KJEE does not prevent someone else from also listening. “Public goods” may be defined as both nonrivalrous and nonexcludable. The extreme corners of these two axes form four named categories and there are many everyday examples for each category (Table 1).

**Table 2.**
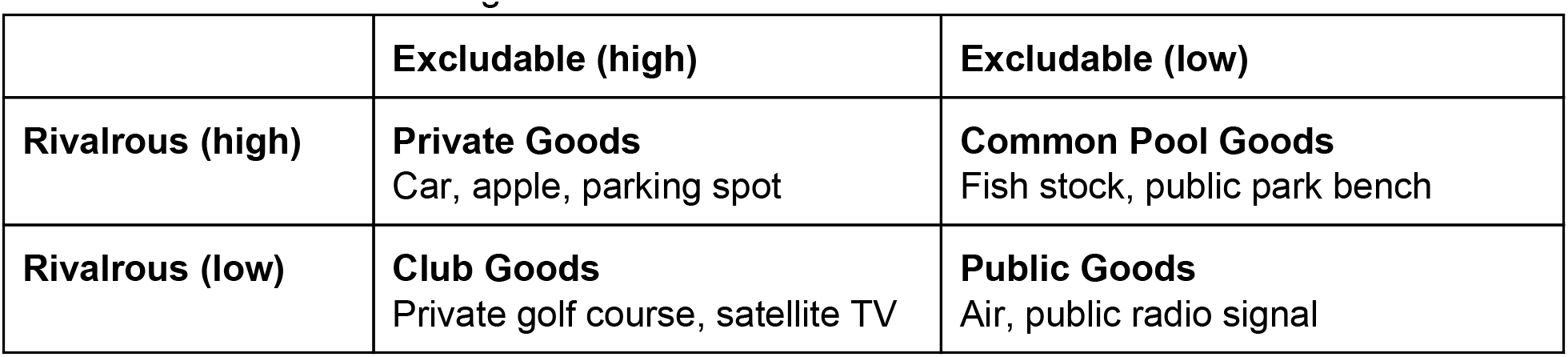
The classification of goods

#### Public Goods in Macroevolution

Recent papers applied the concept of public goods to macroevolutionary topics like the tree of life and novelties. McInerney et al (2011) considered genetic material ("genes” for short) to be a public good and considered the genomes of biological species to be the consumers in this economic metaphor. McInerney et al (2011) contrast their public good hypothesis of genes with the traditional idea of a universally bifurcating ‘tree of life’, with vertical transmission from ancestor to descendant species. With only vertical transmission, genes are highly excludable between species because they can only be present in a genome if inherited from a direct ancestor. In this view, genetic sequences also have low rivalry because no matter how many descendents evolve, they all can have those genes in their genome (assuming use of those genes does not mediate competition between the species). Non-rivalrous, excludable goods of the tree of life model are "club goods”. Instead, McInerney et al (2011) argue that genes should be considered "public goods” because horizontal transfer - the non-vertical transfer of genes to distantly related species - is very common. Horizontal transfer makes genes much less excludable because they could be transferred from any clade to any other clade.

Erwin (2015) applied the concept of public goods to major innovations during evolution. He extends public goods thinking in macroevolution beyond genes to include environmental and ecological goods. In particular, he suggests that the origins of public goods were associated with some major transitions in evolution. A prime example is the production of oxygen, first as a waste product of photosynthetic cyanobacteria. According to Erwin (2015) once oxygen accumulated in the Great Oxygenation Event, it became both non-excludable and non-rivalrous, a public good that was exploited by other organisms.

#### Does an Economic Goods Perspective Advance Macroevolution?

But changing the metaphor alone does not advance the field very far, nor does the frailty of the assumptions of a tree of life metaphor imply that economic goods are the best alternative metaphor. Ideally, an alternative would open doors for new ways of analyzing and understanding macroevolution. Unlike applications of public goods to microevolution, public goods in macroevolution has not matured very far. Macroevolutionary applications differ from public goods applications to microevolution, especially by ignoring costs and by not seeking to optimize anything. Frank (2010) defined a public good as "An individually *costly* act that benefits all group members” [italics added]. The conflict between cost and benefit is central to microevolutionary topics like altruism or cooperation, kin and group selection (Hamilton, 1975), parasite virulence (Frank, 1996), ‘tragedy of the commons’ scenarios, and cases where microbes produce public goods like nitrogen (West et al., 2002) or iron-scavenging molecules (Kümmerli et al., 2009). These topics follow the common practice of economic models that search for equilibria and optima, exploring (for example) whether ‘cheaters’ might exist without bringing down the entire enterprise or whether altruists must exist to maintain the enterprise. Paraphrasing Mr. Spock, sometimes the needs of the many outweigh the needs of the few, or the one (Star Trek, 1982).

Unlike the microevolutionary applications, where a cost is incurred by individuals producing the good or altruistic service, current macroevolutionary ideas do not consider cost. For example, considering oxygen as a public good does not consider that it is an inhibitory waste product for some organisms, yet required by other organisms. Similarly, the Public Goods conception of universally transferrable genes has not investigated equilibria, nor is it directly concerned with costs to the producers. There would seem to be no direct cost to a species if one of its genes is copied into a distant relative. If we posit that competition between genomes is enhanced by the presence of the same genes, then this makes gene function rivalrous, and changes the public good scenario to a pool goods (Table 1) scenario. Perhaps because of this absence of quantification of costs, benefits, or optima, research in phylogenetics has not yet used the mathematical framework of public goods that is present in microevolution, and which might elevate it from metaphor to model. Clearly, there are costs to keeping genes in a genome: they must be copied at each cell division, and their functions could interfere with each other. But phylogenetics is not directly concerned with competition or optima, so perhaps metaphor is as far as economic goods ideas will get. Even if economic models are not the future of macroevolution, these ideas may inspire thinking about the main parameter that separates the tree of life and public goods metaphors, excludability.

#### Excludability

The key difference between the assumptions underlying a strict phylogenetic tree and an economic goods metaphor is excludability. Excludability in economics means use of the product by consumers can be restricted. I propose a particular subset of public goods, the economics of information production (e.g. Benkler, 2002/7; DeLong and Froomkin, 2000), might be an especially appropriate metaphor. DeLong and Froomkin imagined a "New Economy” (DeLong and Froomkin, 1997) where information - non-rival, non-excludable information - is a primary product. Digital media (which now includes scientific publications) provides a useful example for comparing economics to biology because media can easily be copied, making it often non-excludable. At the same time, content owners (like Elsevier) may create copy protection schemes that make digital content more excludable. In fact, the ancient economy of biology may have been trading in non-rival, non-excludable genetic information, eons before the information age of the internet.

I can envision at least two ways in which the concept of excludability is applied to macroevolution. First, we might quantify how excludable biological entities are by examining the distributional patterns of those entities. For example, if genes are shared commonly and homogeneously in all species, even among distant relatives, they are fully non-excludable. This is the idea of McInerney et al (2011) for the species-level tree of life. As they point out, we do not realistically expect every genome to contain all genes, or to be identical, because there are other influences on the content of genomes, like environment or niche. In borrowing from the ecology of microbes, they suggest that every gene is everywhere, but the environment selects (Fondi et al., 2016). They argue fusion (adding genes to the genome of a distant relative) by horizontal transfer is so common, that there is a real possibility to move genes between any two species. From this perspective, we might consider homogeneity of genes across species as a sort of null model. It may be interesting to extend this way of thinking to developmental systems, where we might ask how excludable gene expression is between cell-types or between organs. Is every gene expressed everywhere? In contrast, to what extent do we see gene expression that is exclusive to a particular cell types or organs? Are some cell or tissue types more exclusive than others?

Besides pattern, another way to consider excludability is to consider mechanisms - to identify processes analogous to (for example) the copy protection mechanisms of digital media. At the species level, one such process is reproductive isolation. The Biological Species Concept (BSC) posits that reproductive isolation - the complete prevention of sharing genes outside a well-defined gene pool - is the mechanism to create species. Taking a hard line with this perspective, reproductive isolation makes genomes completely exclusive. The opposite is horizontal transfer mechanisms that make genomes non-excludable. In developmental systems, we have multiple levels of organization to consider. We do not generally expect that animal genes will be horizontally transferred across species and become expressed. However, we do know that within a species, the same genes may come to be expressed in distantly related cell types or organs by co-option. Do the specific mechanisms of co-option lead to different patterns of exclusivity in various cell-types, organs, or morphological features? These are the kinds of questions that an economic goods metaphor can inspire.

### b. Phylogenetic methods

If evolutionary history is treelike, or if we can determine subsets of life’s history that are treelike, we can use the statistical machinery of phylogenetics, developed over decades, to analyze the rising deluge of RNA-seq data and address questions of homology, convergent evolution, novelty, cell-type evolution, and more. If evolution is usually not treelike, we may need a fundamental shift in how we analyze comparative data sets. A first step in data analysis may be to test for "treeness” (Cavalli-Sforza and Piazza, 1975), asking how treelike is a dataset (Kin et al., 2015; Liang et al., 2015). If treelike, then we can proceed with traditional phylogenetic methods. Kin et al. found endometrial stromal cells to have high treeness, generated a cell-type tree for those cells, inferred ancestral states of gene expression, and demonstrated function of reconstructed genes in cell differentiation. Clearly, a fully phylogenetic approach was fruitful. Although they failed to reject a sister-cell null model, there is still evidence that co-option could be important. Namely, 16% of the genes they studied had low values in their treeness metric. Furthermore, they analyzed a similar set of cell types, which could underestimate levels of co-option if particular genes were co-opted to or from cell types they did not sample. They also studied cells from a single species. In multi-species studies, treeness will be increased by species-level history, and new approaches may have to be developed to disentangle species level and cell-type level treeness. Despite my minor critiques, estimating treeness will be important to identify data sets that can be analyzed fruitfully with traditional phylogenetic methods. Of interest will be to determine what types of cell-types or organs, and over what time scales, may be evolving in a more treelike fashion than others.

If datasets reject treelike behavior, we must use phylogenetic network methods. Based especially on the realization of the great extent of horizontal transfer, network models of phylogenetic history are an active area of development, and have been reviewed elsewhere (Huson et al., 2010; e.g. Huson and Bryant, 2006; Strimmer and Moulton, 2000). Compared to pure phylogenetic models, network approaches must model horizontal transfer events, which increases the search space of the problem. Still, the methods have proven useful, especially for understanding the history of genes across genomes. They also should be applicable in some cases to developmental systems (Liang et al., 2015). However, these methods are focused on genetic systems, and assume the units of study are individuated.

A fertile ground for methods development is to understand the history of developmental systems that experienced concerted evolution, fusion, and furcation. Liang et al (2016) quantified concerted evolution of gene expression in vertebrate tissues. Like previous studies (Guo et al., 2007; Gu, 2004; Oakley et al., 2005) Liang et al (2016) assume a Brownian motion model for the evolution of gene expression level. (Note that discretized gene expression data (Wagner et al., 2013) have employed an Mk-like (Lewis, 2001) Markov model, with unified rate parameters across all genes (Oakley et al., 2006) to understand evolution). Liang et al (2016) found rampant concerted evolution, and that morphologically and developmentally related tissues have higher rates of concerted evolution. Another common approach to the evolution of developmental systems is to use multivariate analyses, where each gene is a dimension. Here, studying the evolution of developmental systems will share much in common with phylogenetic multivariate methods, often used for morphological analyses (Adams, 2014a, 2014b; Revell and Collar, 2009). It is an exciting time to be studying the evolution of developmental systems (Dunn et al., 2013). Transcriptomic datasets, down to individual cells (e.g. Macosko et al.,2015), provide the potential, in theory, to quantify all the genes expressed in cells, organs, and tissues. These biological units have undergone complicated evolutionary processes, dividing (furcating), fusing, and individuating to greater or lesser extents. Understanding these processes will require a mixture of phylogenetic, multivariate, and new methods that make the field of developmental systems phylogenetics one to watch for fresh new insights into the macroevolution of complexity, disparity, innovation, and novelty.

## Acknowledgements

Thanks to our lab group, and to C. Simpson, T. Bergstrom, M. Hahn, and B. O’Meara for insightful discussions and comments. Thanks to J. Musser, D. Arendt, and G. Schlosser for organizing the symposium.

